# The Absence of *E. coli* Nucleoid-Associated Protein FIS at Low Temperature Induces an Adaptive Response that Leads To Genome Compaction in Small Rods

**DOI:** 10.1101/2025.08.08.669337

**Authors:** Pamela G. Jones

## Abstract

For *Escherichia coli*, an adaptive response to temperatures just above the minimum temperature of growth, 8°C, includes a change in morphology from rods to small rods. A study was initiated to determine the requirement of nucleoid-associated protein FIS for growth and genome compaction in the small rods at low temperature. Growth and nucleoid staining analyses revealed that the *fis* null mutant displayed decreased growth and initially formed filaments containing decondensed nucleoids at 12°C, indicating that FIS facilitates production of small rods with condensed nucleoids at low temperature. However, characterized by biphasic growth at low temperature, the *fis* null mutant exhibited increased growth, cell division, and nucleoid condensation following a lag phase. Furthermore, compacted circular-shaped nucleoids were formed near the onset of the second growth phase. Therefore *E. coli* responds to the absence of FIS by inducing an adaptive mechanism that causes a shift towards nucleoid condensation resulting in genome compaction in small rods. Furthermore, the deletion of *sulA* (encodes DNA damaged-induced cell division inhibitor SulA) in the *fis* null mutant resulted in suppression of the filamentous morphology. This indicates that the absence of FIS with nucleoid decondensation led to DNA damage, inducing cell division inhibition by SulA.

## INTRODUCTION

A physiological change that occurs upon exposure of *Escherichia coli* to low temperature is an alteration in morphology. While rods are formed at 37°C, small rods are produced at temperatures just above 8°C, the minimum temperature of growth (Porter et al. 2016). The morphological transformation from rods to small rods at low temperature is due to an increase in cell division, specifically requiring late cell division proteins FtsN and DedD (Porter et al. 2016). The formation of the small rod cells is an adaptive response to low temperatures. Mutants that produced small cocci grew like the wild-type strain at 10°C. In contrast, mutants that formed filaments had impaired growth (Porter et al. 2016). Furthermore *E. coli* has been shown to produce small rods and coccoid cells in response to certain other stressful conditions in order to conserve energy and to optimize the influx of nutrients and efflux of wastes (Young, 2007).

The volume of ∼1.6 mm *E. coli* chromosomal DNA must effectively reduce to sufficiently fit within the cell. In addition to macromolecular crowding and supercoiling, nucleoid-associated proteins contribute to DNA condensation (Luijusterburg et al. 2006; Luijusterburg et al. 2008). Serving an architectural role on the chromatin, nucleoid-associated proteins are small basic proteins that can wrap, bend or bridge distant DNA segments (Luijusterburg et al. 2006; Luijusterburg et al. 2008). One of several nucleoid-associated proteins in *E. coli*, FIS can specifically recognize a 15-bp motif as well as nonspecifically bind to DNA (Pan et al. 1996; Shao et al. 2008; Stella et al. 2010). Magnetic tweezers experiments showed that FIS binding to non-specific sites resulted in significant DNA compaction due to DNA bending as well as formation and stabilization of DNA loops (Skoko et al. 2005; Skoko et al. 2006). FIS is the most abundant chromatin protein during exponential growth (Talukder et al. 1999). Although the *fis* null mutant displayed a longer cell length and dispersed nucleoids at 30°C (Lioy et al. 2018), a specific growth requirement of FIS for global compaction has not been clearly demonstrated.

Because a role for FIS in determining nucleoid and cellular morphology at low temperature had not been investigated, a study was initiated to determine the requirement of FIS for growth, nucleoid condensation, and small rod morphology at temperatures near the minimum temperature of growth. In this work, the data indicate that FIS promotes growth and facilitates the organization of compacted nucleoids in the small rods. The *fis* null mutant exhibited biphasic growth accompanied by transient production of filaments containing large decondensed nucleoids at 12°C. Furthermore, the data demonstrate that *E. coli* responds to the lack of FIS by inducing an adaptive mechanism that leads to the formation of small rods with condensed nucleoids. Extragenic suppressor mutations of the *fis* null mutation are also identified that led to an increase in nucleoid condensation and/or cell division.

## MATERIALS AND METHODS

### Bacterial strains, plasmids, and growth conditions

The bacterial strains utilized in this study were *E. coli* wild-type strain BW25113 and the corresponding Keio deletion mutants JW3229 (Δ*fis*), JW5741 (Δ*rnr*), JW0941 (Δ*sulA*), JW1935 (Δ*rcsA*) and JW0460 (Δ*ybaB*) (Baba et al. 2006). From dilution of overnight cultures, growth of cultures in Luria-Bertani (LB) media at 37°C was assayed spectrophotometrically at absorbance 420 nm. At an optical density 420 nm of 0.5 to 0.7, the cultures were shifted to 10°C or 12°C followed by overnight incubation. After dilution of the cultures in fresh LB media, growth at low temperature was monitored at absorbance 420 nm. When provided, ampicillin at 50 μg/ml or kanamycin at 30 μg/ml was administered to the cultures. Plasmid pKD123 encoding cell division protein FtsN (Dai et al. 1993) and plasmid pKV1238-*hsIVU* encoding two component protease HsIVU (Kanemori et al. 1999) were utilized in this study.

### Construction of double null mutants

As a result of transformation into *fis::kan* mutant strain JW3229, plasmid pCP20 (Cherepanov et al. 1995) containing a temperature-sensitive origin of replication was used to remove the kanamycin resistance gene generating an ampicillin sensitive and kanamycin sensitive *fis* deletion strain. P1 transduction was used to transfer the *rnr::kan* mutation in JW5741 or *sulA::kan* mutation in JW0941 or *rcsA::kan* mutation in JW1935 or *ybaB::kan* mutation in JW0460 into the kanamycin sensitive *fis* deletion mutant. The double mutants were isolated by growth on LB plates containing kanamycin at 37°C. Polymerase chain reactions were done to confirm the presence of the null mutations.

### Microscopy & Staining

Heat-fixed cells were stained with 0.35% crystal violet, and cellular morphology was examined by light microscopy using BS41 brightfield microscope under a total magnification of 1,000X. Photographic images were taken using a DP71 microscope digital camera. Ethanol-fixed cells were applied to poly-L-lysine coated slides and were stained with DAPI (4′,6-diamidino-2-phenylindole) in slow fade gold antifade reagent (Invitrogen) as described (Usongo et al. 2008). Stained cells were viewed with Olympus BX43 equipped with 100 W mercury lamp and DAPI filters under a total magnification of 1,000X. Phase contrast optical system of Olympus BX43 was used to view cells under a total magnification of 1,000X. Photographs were taken with a DP23M digital microscope camera, and images were analyzed using Cellsens software version 4.2.

## RESULTS

### FIS facilitates growth and the small rod morphology at low temperature

Because cells lacking FIS grow similarly as the wild-type but are elongated at 30°C (Lioy et al. 2018), experiments were done to determine whether FIS is required for growth and the small rod morphology at low temperature. In comparison to growth of the wild-type strain BW25113, the growth of *fis* null mutant was reduced at 10°C (Fig 1). Exhibiting biphasic growth, the *fis* null mutant underwent a lag or acclimation phase beginning at the 72 hr followed by increased growth at the 168 hr. The growth of the *fis* null mutant was slower at the second growth phase. The generation time at the first phase was 28 hr compared to 67 hr at the second phase. In comparison, the generation time of the wild-type strain BW25113 was 18 hr. Microscopic examination showed that the wild-type strain BW25113 formed rods with an average size of 2.5 μm (Fig. 2A) and the *fis* null mutant formed elongated rods with an average size of 3.6 μm (Fig. 2B) at 37°C. In comparison to the rods formed at 37°C (Fig. 2A), wild-type strain BW25113 produced small rods at 10°C at the 48 hr (Fig 2C). In contrast, the *fis* null mutant formed filaments at the 48 hr during the first phase of growth (Fig. 2D) and at the end of the lag phase at the 168 hr (Fig. 2E). The average size of the filament is 9.1 μm compared to the average size of 1.0 μm of the small rods of the wild-type strain BW25113. However at the 264 hr during the second phase of growth, the *fis* null mutant produced small rods (Fig. 2F). Therefore, the function of FIS leads to an increase in cell division resulting in growth and the small rod morphology low temperature. Furthermore, the biphasic growth of the *fis* null mutant and production of small rods in the second growth phase indicate that *E. coli* acquired a mechanism to adapt to the absence of FIS at low temperature.

**Fig 1.**
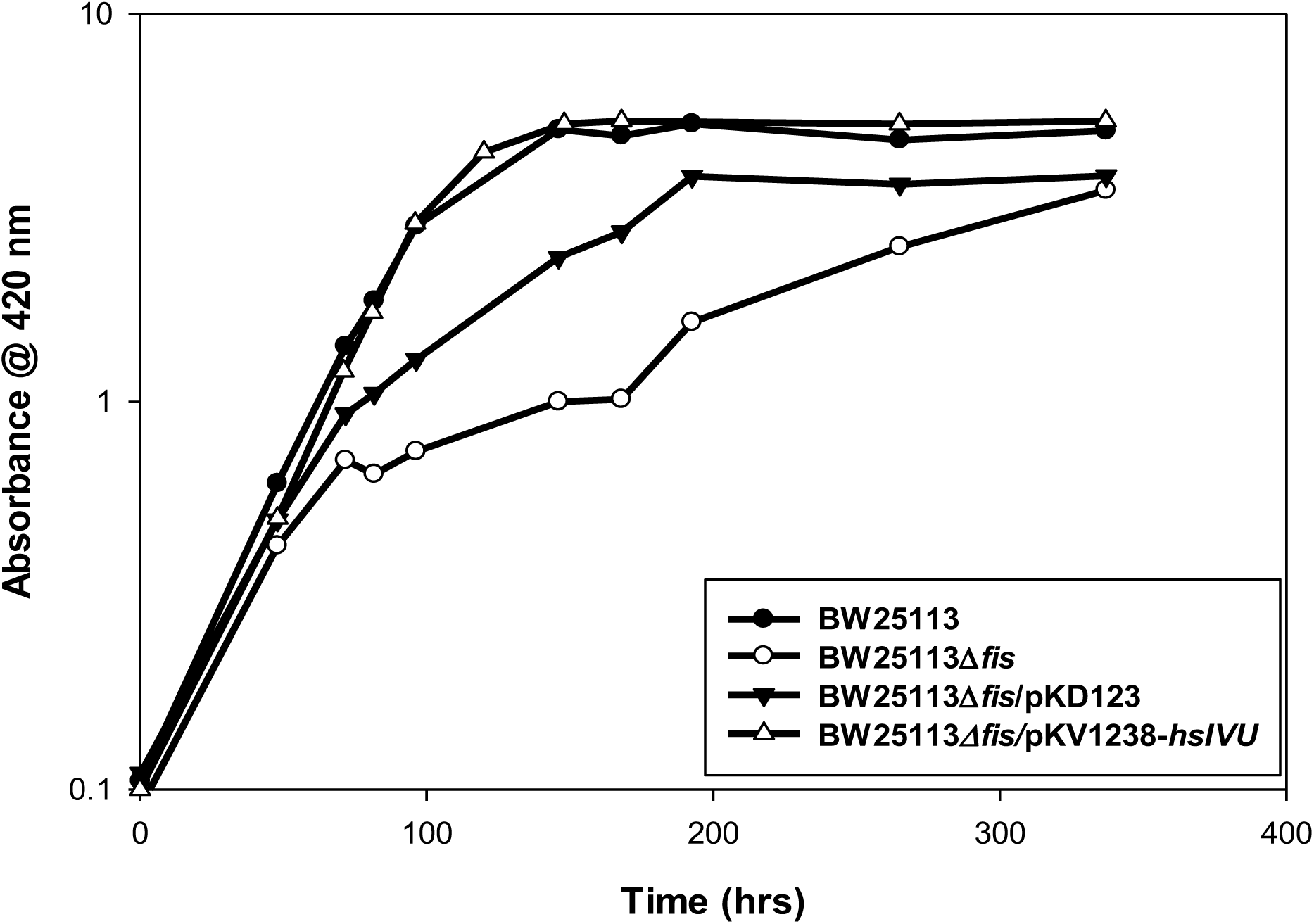
Effect of FIS on growth at 10°C. Growth of the wild-type strain BW25113, *fis* null mutant, and *fis* null mutant containing plasmid pKD123 encoding cell division protein FtsN and containing plasmid pKV1238-*hsIVU* encoding two component protease HsIVU at 10°C. Exponentially growing cultures in LB media at 37^ο^C were shifted to 10°C. After overnight incubation, the strains were diluted in LB media, and growth was monitored spectrophotometrically at absorbance 420 nm for the times indicated

**Fig. 2.**
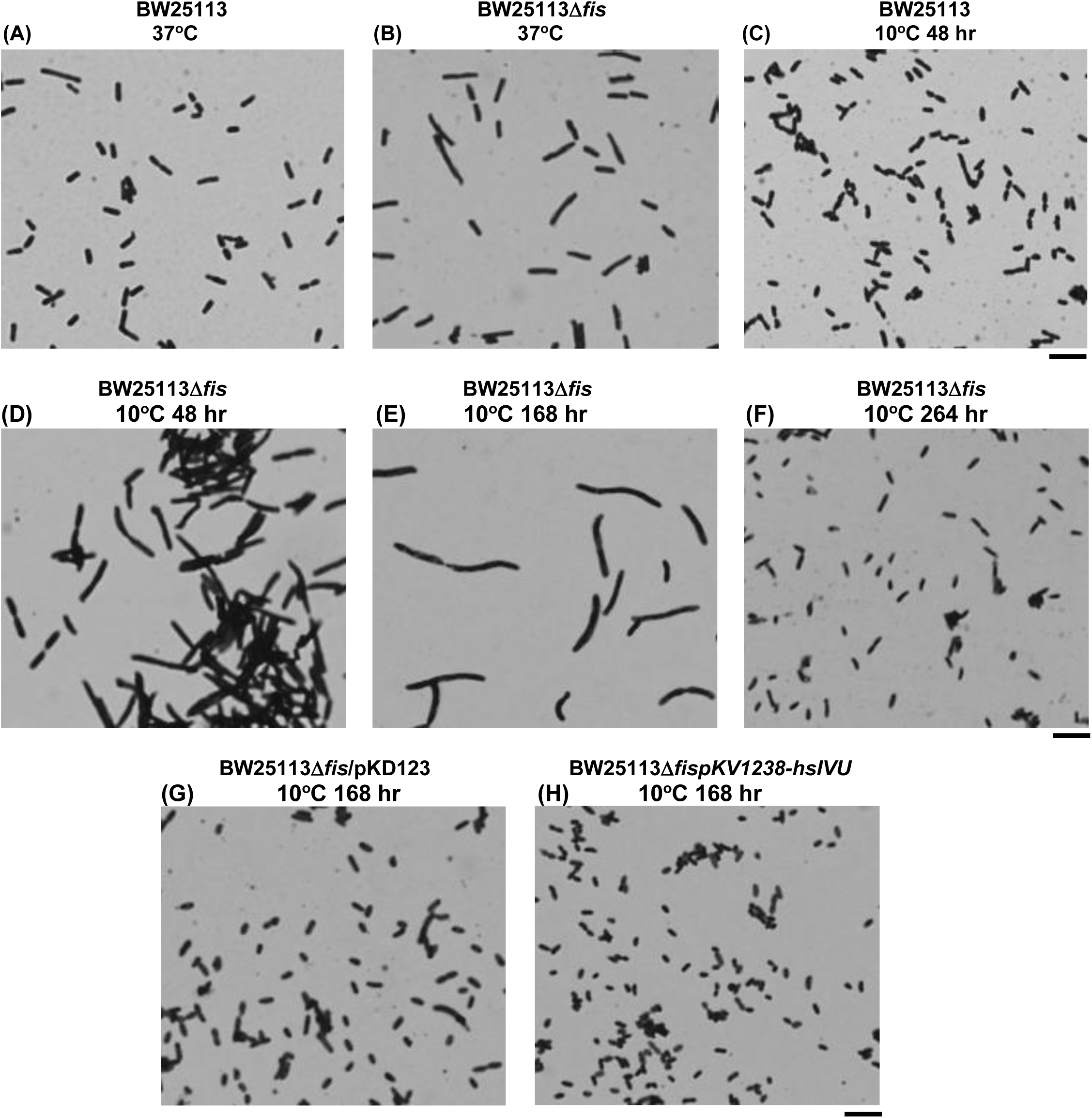
Effect of FIS on cellular morphology at 10°C. Cellular morphology of the wild-type strain BW25113 and *fis* null mutant at 37°C (A & B). Cellular morphology of the wild-type strain BW25113, *fis* null mutant, and *fis* null mutant containing plasmid pKD123 encoding cell division protein FtsN as well as containing plasmid pKV1238-*hsIVU* encoding the two component protease HsIVU at 10°C (C-H) at the times indicated. The cells were stained with crystal violet. (Bar = 5 μm)

### hslVU is a multicopy suppressor of the fis null mutation

Cell division protein FtsN is absolutely required to increase septation resulting in the formation of small rods at low temperature (Porter et al. 2016). Experiments were done to determine whether *ftsN* is a multicopy suppressor of the cold-sensitive phenotypes of the *fis* null mutant. Plasmid pKD123 encodes FtsN (Dai et al. 1993). The presence of plasmid pKD123 in the *fis* null mutant resulted in increased growth (Fig. 1). In comparison to the filaments formed by the *fis* null mutant at the 168 hr at 10°C (Fig. 2E), the presence of the plasmid pKD123 in the mutant resulted in the formation of small rods (Fig. 2G). However, the presence of other FtsN-encoding plasmids in the *fis* null mutant failed to restore the wild type phenotypes (data not shown). Subsequent analysis of the plasmid pKD123 revealed that, in addition to gene *ftsN*, genes *hsIV* and *hslU* could also be expressed by the plasmid. Located adjacent to *ftsN*, genes *hslV* and *hslU* encoding HsIV (ClpQ) and HslU (ClpY), respectively, comprise the two-component protease HslVU (Rohrwild et al. 1997). HslU is the catalytic subunit and HslV is the ATPase subunit. Experiments were conducted to determine whether *hslVU*, rather than *ftsN*, is the multicopy suppressor of the cold-sensitive phenotypes of the *fis* null mutant. Plasmid pKV1238-*hslVU* encodes the HslVU protease (Kanemori et al. 1999). In comparison to the impaired growth of the *fis* null mutant at 10°C, the presence of plasmid pKV1238-*hslVU* in the mutant resulted in growth as similarly observed as the wild-type strain BW25113 (Fig. 1). In contrast to the filaments formed by the *fis* null mutant at the 168 hr at 10°C (Fig. 2E), the presence of plasmid pKV1238-*hslVU* in the mutant resulted in the formation of small rods (Fig. 2H). The data indicate that *hslVU i*s a multicopy suppressor of the cold-sensitive phenotypes of the *fis* null mutant.

### FIS facilitates genome compaction at low temperature

Because the *fis* null mutant exhibited biphasic growth at 10°C (Fig. 1), the growth pattern of the mutant was also determined at 12°C. As shown in Fig 3, the *fis* null mutant displayed biphasic growth. The first phase of growth with a generation time of 27 hr was followed by a lag phase and a subsequent second phase of growth with a generation time of 31 hr. In comparison, the generation time of the wild-type strain BW25113 was 15 hr. At 12°C, the lag phase was shorter compared to 10°C. In contrast to 96 hours at 10°C, the lag phase at 12°C was 20 hours, beginning at the 72 hr and ending at the 92 hr. Therefore, it takes longer for *E. coli* to adapt to the absence of FIS at 10°C compared to 12°C.

**Fig. 3.**
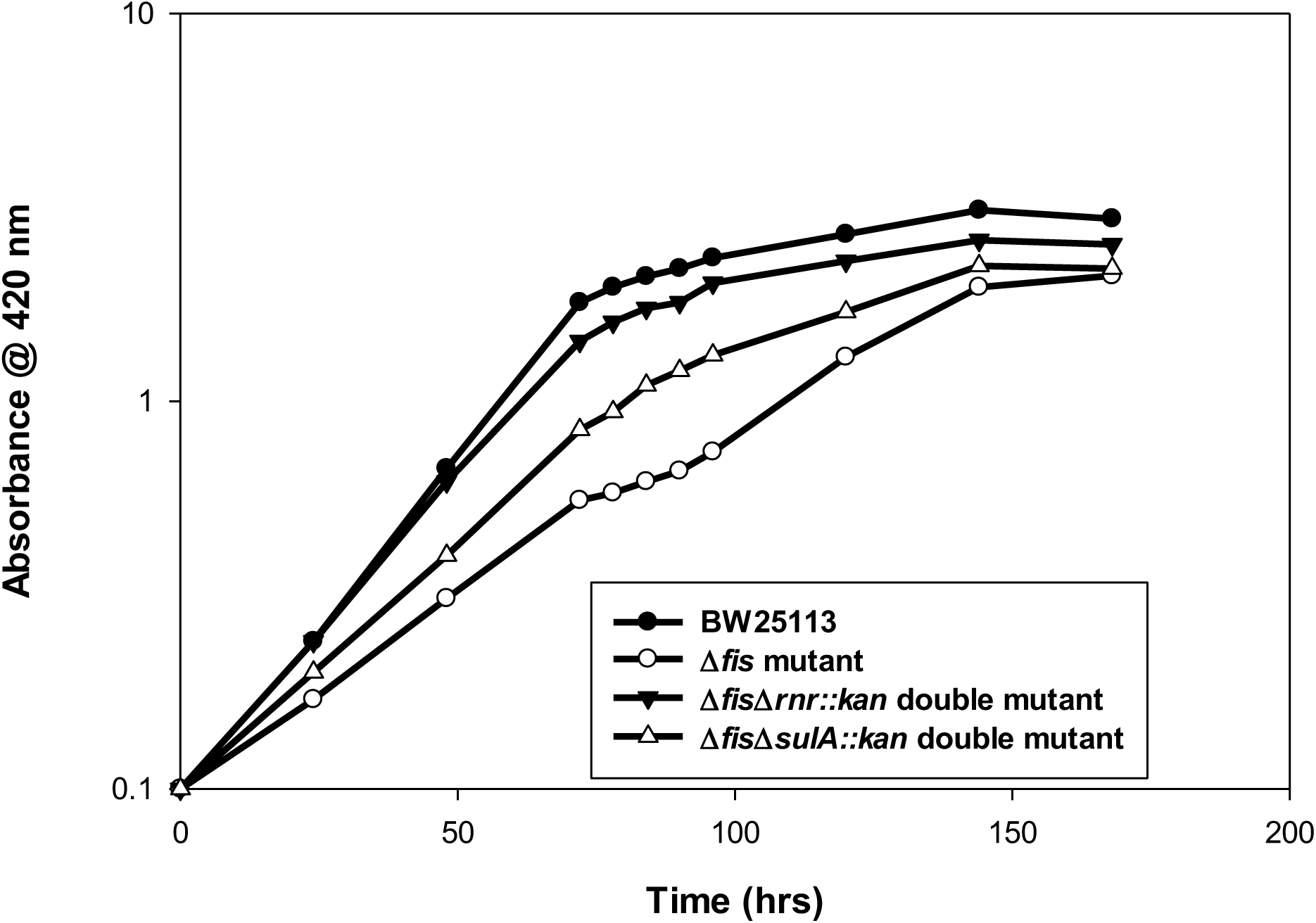
Effect of FIS on growth at 12°C. Growth of the wild-type strain BW25113, the Δ*fis* mutant, Δ*fis*Δ*rnr::kan* double mutant and Δ*fis*Δ*sulA::kan* double mutant at 12°C. Exponentially growing cultures in LB media at 37^ο^C were shifted to 12°C. After overnight incubation, the strains were diluted in LB media, and growth was monitored spectrophotometrically at absorbance 420 nm for the times indicated

DAPI staining of the nucleoid was done to examine the effect of FIS on nucleoid structure at 12°C. Phase-contrast and fluorescence microscopy was used for viewing of cellular and nucleoid morphology. In contrast to the compacted nucleoids located in small rods of the wild-type strain BW25113 (Fig. 4 A-C), the nucleoids located in the Δ*fis* mutant cells were decompacted throughout the filaments (Fig. 4 D-F) at the 72 hr, which is the start of the lag phase. The average length of the filaments is 6.2 μm compared to the 1.1 μm average size of the small rods of the wild-type strain BW25113. At the 96 hr at the second phase of growth, the *fis* null mutant displayed filaments but also shorter filaments and rods that were formed as a result of an increase in cell division (Fig. 5C). Cell constrictions leading to formation of shorter cells are also observed, which is consistent with the increase in growth at the second phase. In addition, DAPI staining revealed an uneven distribution of the nucleoid (Fig. 5 A & B). Furthermore as shown in Fig. 5 A & B, there was an increase in nucleoid condensation. The microscopic images showed cells with circular compacted nucleoids, including some nucleoids that appear closely packed or merged together (indicated by the red arrows in Fig. 5B). Enlarged images of the cells with the circular compacted and closely packed or merged nucleoids are shown in Fig. 5 D-F. Red arrows point to the nucleoids in Fig. 5E. By the 120 hr, there was a continued increase in cell division accompanied with genome compaction in the small rods (Fig. G-I). The average size of the small rods of the *fis* null mutant at the120 hr is 1.4 μm compared to 1.1 μm average size of the small rods of wild-type strain BW25113 and 6.2 μm average size of the filaments of the *fis* null mutants at the 72 hr. Consistent with uneven nucleoid distribution observed at the 96 hr, anucleate cells were also formed (Fig. 5G). The data demonstrate that FIS facilitates production of small rods with condensed nucleoids. However, the absence of FIS with nucleoid decondensation induces an adaptive response that facilitates growth accompanied by an increase in cell division and genome compaction.

**Fig. 4.**
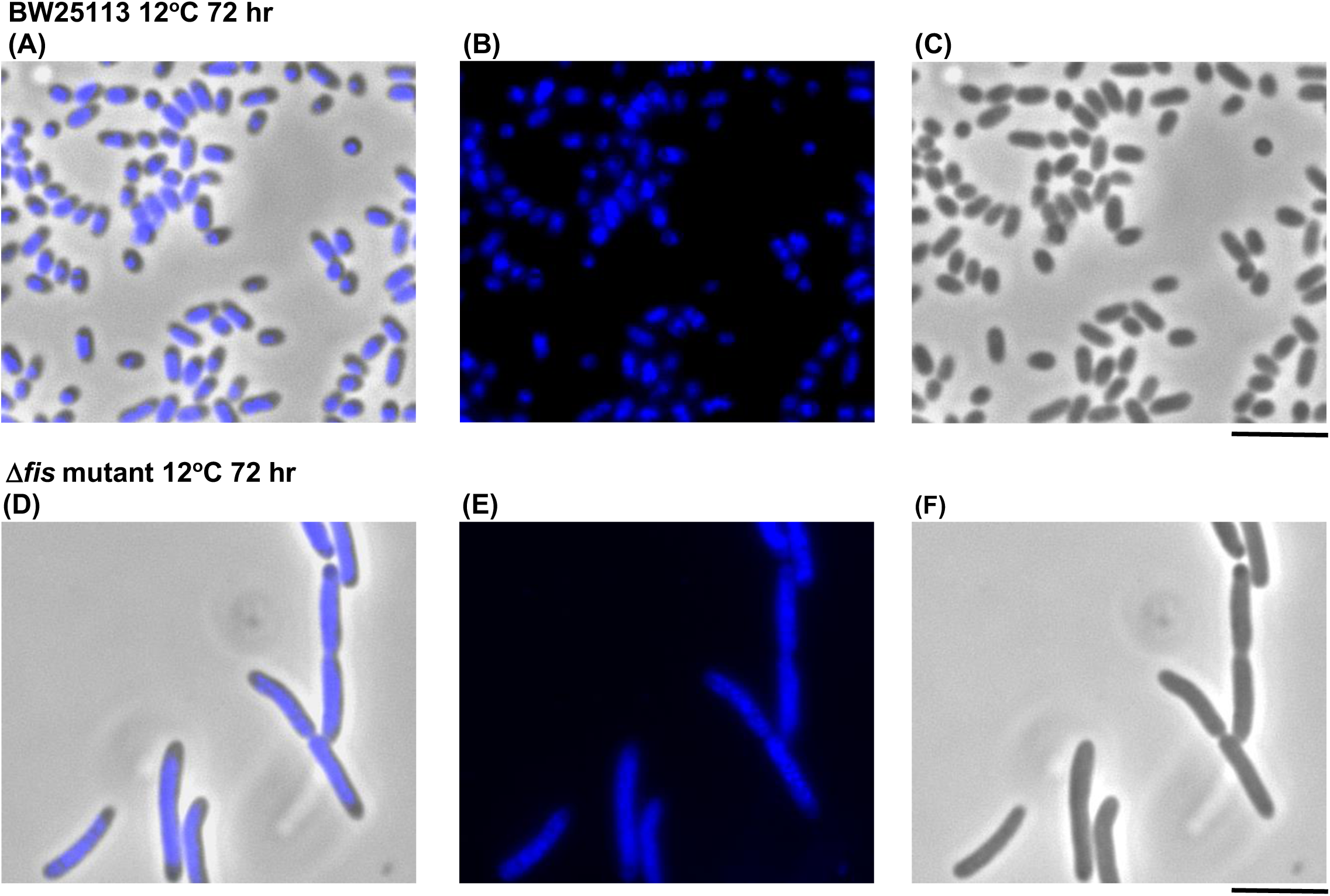
Effect of FIS on cellular and nucleoid morphology at 12°C. Phase-contrast and fluorescent DAPI-stained (false-blue) overlay images (A & D), fluorescent DAPI-stained (false-blue) images (B and E), and phase-contrast images (C & F) of wild-type strain BW25113 and the Δ*fis* mutant at 12°C at 72 hr. (Bar = 5 μm)

**Fig. 5.**
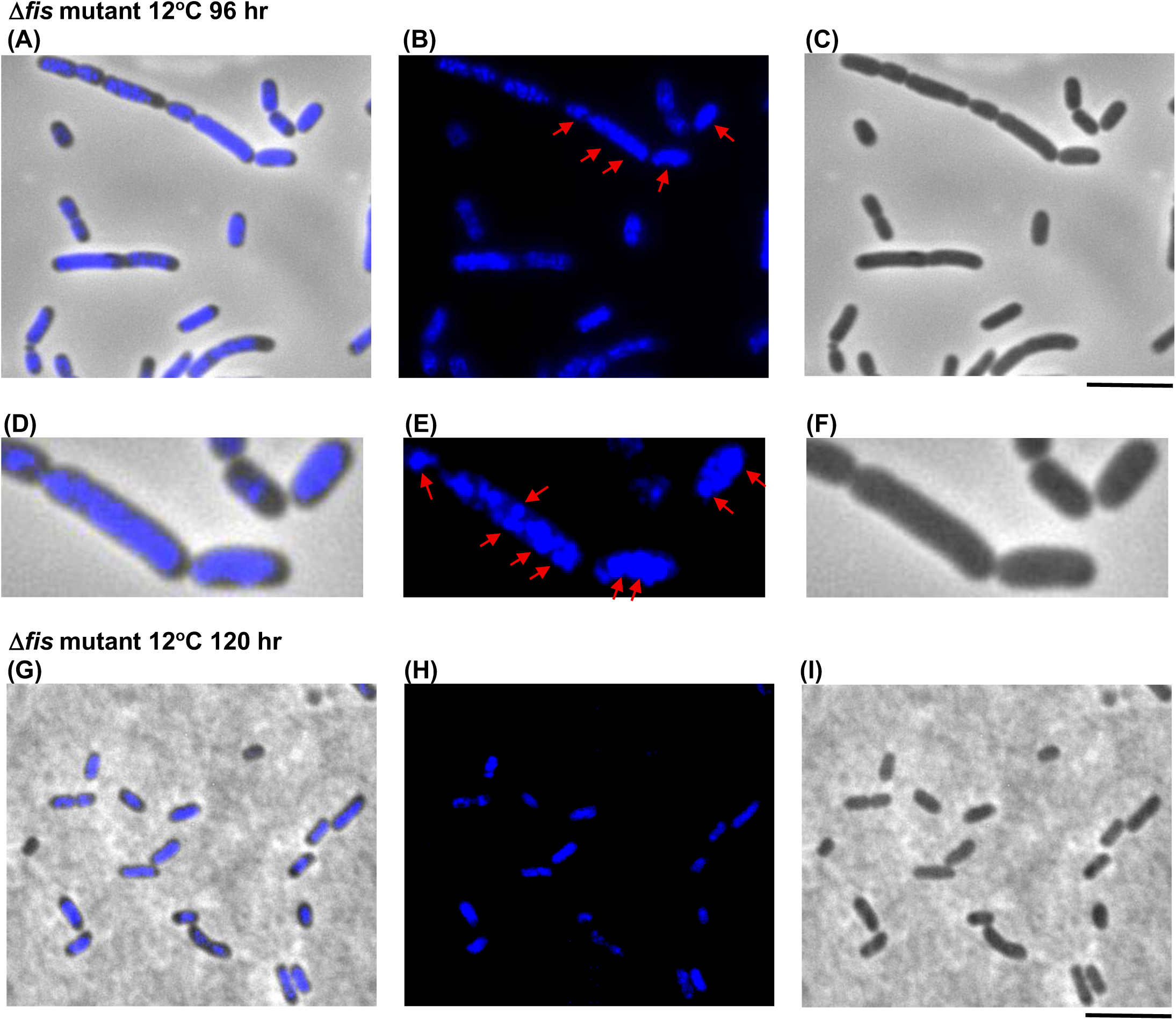
Cellular and nucleoid morphology of Δ*fis* mutant at 12°C at the 96 hr and 120 hr. Phase-contrast and fluorescent DAPI-stained overlay images (A & D), fluorescent DAPI-stained images (B & E), and phase-contrast images (C & F) of Δ*fis* mutant at 12°C at 96 hr. Enlarged images of cells with circular compacted nucleoids shown in D-F. Red arrows point to circular compacted nucleoids. Phase-contrast and fluorescent DAPI-stained overlay image (G), fluorescent DAPI-stained image (H), and phase-contrast image (I) of the Δ*fis* mutant at 12°C at 120 hr. (Bar = 5 μm)

### Null mutations of rnr and sulA are extragenic suppressors of fis null mutation

The finding that *hslVU* is a multicopy suppressor of the cold-sensitive phenotypes of the *fis* null mutant implies that specific destabilization of a substrate by the HsIVU protease leads to alleviation of the functional requirement of FIS at low temperature. Natural protein substrates of the HsIVU (ClpQY) protease include SOS-induced cell division inhibitor SulA, capsule synthesis activation protein RcsA, 3’-5’ exoribonuclease RNase R, and nucleoid-associated protein YbaB (Tsai et al. 2017). Therefore, experiments were done to determine whether a null mutation of *sulA* or *rcsA* or *rnr* or *ybaB* is an extragenic suppressor of the *fis* null mutation at low temperature. Growth and microscopic analyses of wild-type strain BW25113, Δ*fis* mutant, and the various double mutants were performed at 12°C. The deletion of *rcsA* or *ybaA* from the *fis* null mutant did not result in improved growth nor the formation of small rods (data not shown). However in contrast to the reduced growth of the Δ*fis* mutant, the Δ*fis*Δ*rnr::kan* double mutant grew similarly as the wild-type strain BW25113 (Fig. 3). Furthermore, the presence of the *rnr* null mutation suppressed the aberrant filamentous and nucleoid morphology of the Δ*fis* mutant resulting in the increased production of small rods with compacted nucleoids at the 72 hr (Fig. 6 A-C). The average size of the Δ*fis*Δ*rnr::kan* double mutant cells is 1.6 μm. A cell containing circular condensed nucleoids with two nucleoids that appear merged together is also observed as indicated by the red arrows in Fig. 6B. Enlarged images of the cells with the circular condensed nucleoids are shown in Fig. 6 D-F. Red arrows point to the nucleoids in Fig. 6E. The data indicate that *rnr* null mutation is an extragenic suppressor of the cold-sensitive phenotypes of the *fis* null mutant. In comparison, the *sulA* null mutation is a partial suppressor of the cold-sensitive phenotypes of the *fis* null mutant. The deletion of the *sulA* gene from the *fis* null mutant led to improved growth (Fig. 3) and increased production of rods, with an average size of 2.9 μm, mainly containing dispersed nucleoids at the 72 hr (Fig. 6 G-I).

**Fig. 6.**
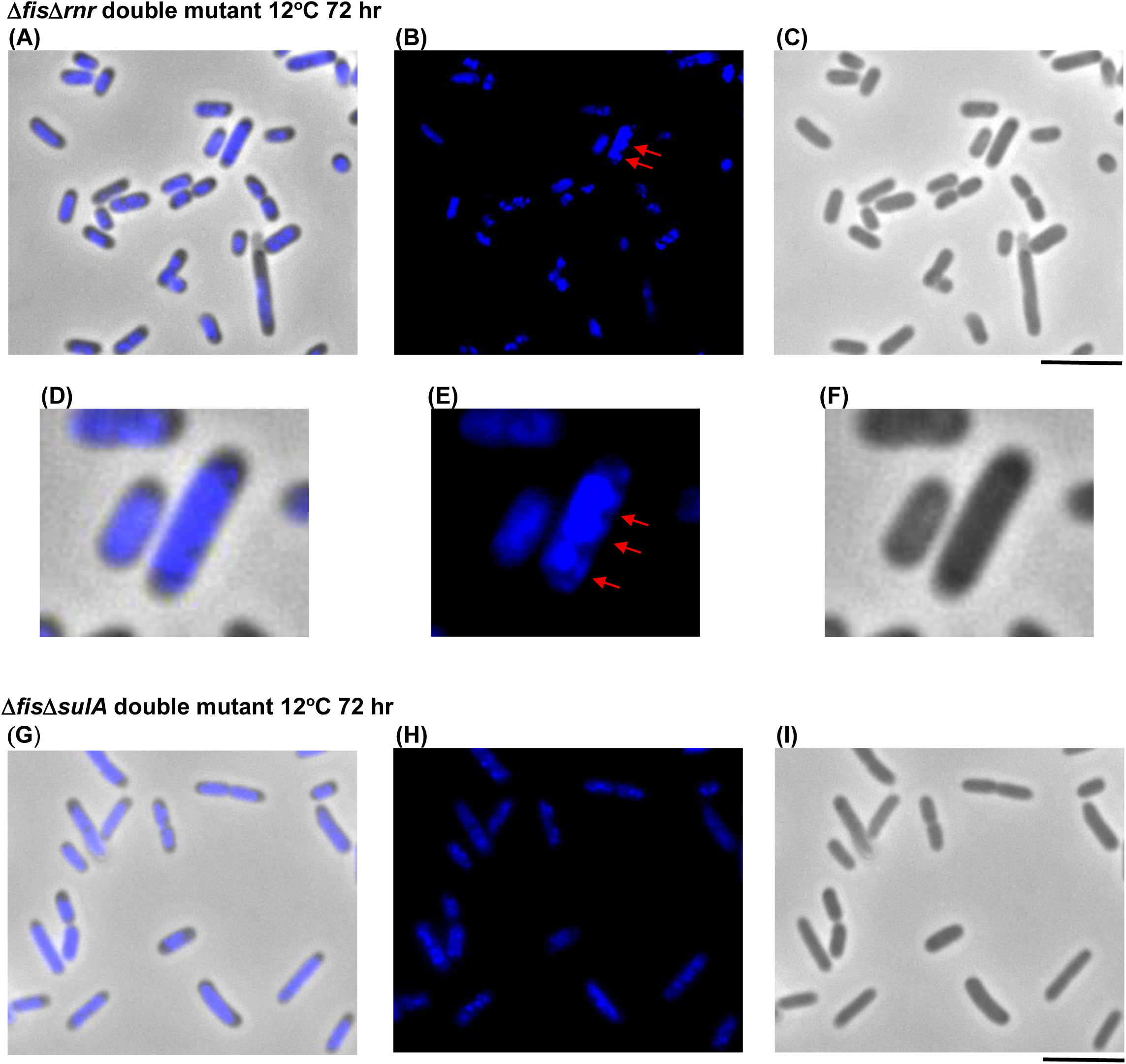
Extragenic suppression of aberrant nucleoid and cell morphology of the Δ*fis* mutant. Phase-contrast and fluorescent DAPI-stained overlay image (A & D), fluorescent DAPI-stained image (B & E), and phase-contrast image (C & F) of the Δ*fis*Δ*rnr::kan* double mutant at 12°C at 72 hr. Enlarged images of cells with circular compacted nucleoids are shown in D-F. Red arrows point to circular compacted nucleoids. Phase-contrast and fluorescent DAPI-stained overlay image (G), fluorescent DAPI-stained image (H), and phase-contrast image (I) of the Δ*fis*Δ*sulA::kan* double mutant at 12°C at 72 hr. (Bar = 5 μm)

## DISCUSSION

A DNA binding and bending protein, FIS serves an architectural role on the chromatin. Although *in vitro* studies showed that FIS DNA binding to nonspecific sites resulted in substantial compaction due to DNA bending as well as formation and stabilization of DNA loops (Skoko et al. 2005; Skoko et al. 2006), an *in vivo* growth requirement of FIS for genome compaction has not been clearly demonstrated. In this work, we examined the requisite of nucleoid-associated protein FIS for growth, nucleoid condensation, and small rod morphology at low temperature. Unlike growth and small rods with condensed nucleoids phenotypic of the wild-type at 12°C, the *fis* null mutant exhibited reduced growth and initially produced filaments with decondensed nucleoids. The data indicate that FIS promotes growth and facilitates formation of small rods with compacted genomes at low temperature.

Biphasic growth typically describes the growth pattern of bacteria in response to the presence of two sugars. However, unlike at high temperature, the *fis* null mutant exhibited biphasic growth at low temperature. This finding indicates that the absence of FIS with nucleoid decondensation is a physiologically stressful condition at low temperature that induces an adaptive response termed the FIS null adaptation response. As a result, multiple and simultaneous cell divisions accompanied by nucleoid condensation occur leading to the production of small rods with compacted nucleoids at the second growth phase. Furthermore, there is a shift towards nucleoid condensation early in the second growth phase as evident by the formation of compacted circular nucleoids, some which are merged or closely packed together. These compacted circular and merged nucleoids resemble the condensed spherical and fused nucleoids induced by chloramphenicol as a result of an increase in nucleoid condensation (van Helvoort et al. 1996; Zusman et al. 1973). However in contrast to the condensed nucleoids induced by chloramphenicol treatment, the compact nucleoids physiologically induced by the lack of FIS with nucleoid decondensation is part of an adaptive response resulting in genome compaction in small rods at low temperature.

In *E. coli*, adaptation to temperatures just above the minimum temperature of growth, 8°C, requires the morphological transformation from rods to small rods (Porter et al. 2016). Because the cells are small and space for the nucleoid is limited, the organization of highly condensed nucleoids is particularly advantageous at low temperature. Therefore, production of small rods with densely compacted nucleoids is beneficial for growth at low temperatures. This is corroborated by the observation that expression of a mutant nucleoid-associated protein HU in *E. coli* resulted in small coccoid cells with tightly condensed nucleoids accompanied with a markedly higher growth rate particularly at low temperatures (Kar et al. 2005). Therefore, FIS mediates an adaptive link between nucleoid condensation and small rod morphology facilitating the compaction of nucleoids in the small rods for growth at low temperature.

The finding that *hslVU* is a multicopy suppressor of the cold-sensitive phenotypes of the *fis* null mutant indicates that destabilization of a substrate by the HsIVU protease leads to the alleviation of the requirement of FIS at low temperature. RNase R is a natural substrate of the HsIVU protease (Tsai et al. 2017). The data showed that a *rnr* null mutation is an extragenic suppressor of the abnormal growth as well as the aberrant nucleoid and cellular morphology of the *fis* null mutant. This implies that proteolysis of RNase R by the HsIVU protease suppresses the functional requirement of FIS to facilitate genome compaction in the small rods. RNase R, a 3’-5’ exoribonuclease, functions in the maturation and decay of highly structured RNAs (Andrade et al. 2009). RNase R also plays a role in regulating gene expression at low temperature (Phadtare 1996). In addition to the upregulation of several genes, DNA microarray analysis revealed that several genes are downregulated in the *rnr* null mutant, including *sulA* encoding the SOS-induced cell division inhibitor (Mukherjee et al 1996; Phadtare 1996). This suggests that the reduced expression of *sulA* contributes to the increase in cell division observed in the *ΔfisΔrnr::kan* double mutant. This is consistent with the finding that the cell division block in the *fis* null mutant is dependent upon SulA. In addition, genes encoding several membrane proteins are downregulated in the *rnr* null mutant at low temperature (Phadtare 1996). Transertion, which is coupled transcription and translation with translocation of nascent membrane proteins, is proposed to promote nucleoid expansion by facilitating attachment of the nucleoid to the membrane by transcription-translation-translocation chain (Woldringh et al. 1995; Woldringh 2002). Cumulative data from various studies indicate that chloramphenicol inhibits nucleoid expansion by causing membrane detachment of the nucleoid leading to nucleoid condensation and fusion (Bakshi et al. 2014; Bakshi et al. 2015; Span et al. 2025). The data showed that the presence of *rnr* null mutation in the *fis* null mutant resulted in nucleoid compaction. It is plausible that reduced co-translation translocation of the membrane proteins (as consequence of reduced expression due to the absence of RNaseR) leads to membrane detachment of the nucleoid resulting in nucleoid condensation in the *ΔfisΔrnr::kan* double mutant. This is supported by the observation of a *ΔfisΔrnr::kan* double mutant cell with circular compacted nucleoids, which bear a resemblance to the spherical condensed nucleoids induced by chloramphenicol treatment (van Helvoort et al. 1996; Zusman et al. 1973). The data suggest that the absence of RNase R in the *fis* null mutant effects gene expression, leading to a bypass of the requirement of FIS for formation of small rods with condensed nucleoids at low temperature.

Cell division inhibitor SulA, which is induced by DNA damage as part of the SOS response (Mukherjee et al; 1998), is a natural substrate of the HsIVU protease (Tsai et al. 2017). The data showed that a *sulA* null mutation is an extragenic suppressor of the filamentous phenotype of the *fis* null mutant at low temperature resulting in the formation of rods. However, the *sulA* mutation did not restore nucleoid compaction, which is consistent with the partial growth restoration of the *fis* null mutant. The suppression of the filamentous phenotype as a result of the *sulA* deletion indicates that the absence of FIS at low temperature induces cell division inhibition by SulA. A generic response to stress, nucleoid compaction serves to insulate the DNA (Shechter et al. 2013). Under stressful conditions, an additional role for chromatin binding of nucleoid-associated proteins is to avert damage (Hołówka & Zakrzewska-Czerwińska 2020). The data indicate that the lack of FIS with nucleoid decompaction causes DNA to be susceptible to damage, triggering a SulA-block in cell division. Therefore, FIS-mediated nucleoid compaction aids to preserve genome integrity at low temperature.

## Acknowledgements

The author thanks The Coli Genetic Stock Center for providing plasmids pKD123 and pCP20 as well as null mutants of the Keio collection. The author also expresses gratitude to Dr. Masaaki Kanemori for kindly providing plasmid pKV1238-*hsIVU*.

